# The Genomic History Of Southeastern Europe

**DOI:** 10.1101/135616

**Authors:** Iain Mathieson, Songül Alpaslan Roodenberg, Cosimo Posth, Anna Szécsényi-Nagy, Nadin Rohland, Swapan Mallick, Iñigo Olalde, Nasreen Broomandkhoshbacht, Francesca Candilio, Olivia Cheronet, Daniel Fernandes, Matthew Ferry, Beatriz Gamarra, Gloria González Fortes, Wolfgang Haak, Eadaoin Harney, Eppie Jones, Denise Keating, Ben Krause-Kyora, Isil Kucukkalipci, Megan Michel, Alissa Mittnik, Kathrin Nägele, Mario Novak, Jonas Oppenheimer, Nick Patterson, Saskia Pfrengle, Kendra Sirak, Kristin Stewardson, Stefania Vai, Stefan Alexandrov, Kurt W. Alt, Radian Andreescu, Dragana Antonović, Abigail Ash, Nadezhda Atanassova, Krum Bacvarov, Mende Balázs Gusztáv, Hervé Bocherens, Michael Bolus, Adina Boroneanţ, Yavor Boyadzhiev, Alicja Budnik, Josip Burmaz, Stefan Chohadzhiev, Nicholas J. Conard, Richard Cottiaux, Maja Čuka, Christophe Cupillard, Dorothée G. Drucker, Nedko Elenski, Michael Francken, Borislava Galabova, Georgi Ganetovski, Bernard Gély, Tamás Hajdu, Veneta Handzhyiska, Katerina Harvati, Thomas Higham, Stanislav Iliev, Ivor Janković, Ivor Karavanić, Douglas J. Kennett, Darko Komšo, Alexandra Kozak, Damian Labuda, Martina Lari, Catalin Lazar, Maleen Leppek, Krassimir Leshtakov, Domenico Lo Vetro, Dženi Los, Ivaylo Lozanov, Maria Malina, Fabio Martini, Kath McSweeney, Harald Meller, Marko Menđušić, Pavel Mirea, Vyacheslav Moiseyev, Vanya Petrova, T. Douglas Price, Angela Simalcsik, Luca Sineo, Mario Šlaus, Vladimir Slavchev, Petar Stanev, Andrej Starović, Tamás Szeniczey, Sahra Talamo, Maria Teschler-Nicola, Corinne Thevenet, Ivan Valchev, Frédérique Valentin, Sergey Vasilyev, Fanica Veljanovska, Svetlana Venelinova, Elizaveta Veselovskaya, Bence Viola, Cristian Virag, Joško Zaninović, Steve Zäuner, Philipp W. Stockhammer, Giulio Catalano, Raiko Krauß, David Caramelli, Gunita Zariņa, Bisserka Gaydarska, Malcolm Lillie, Alexey G. Nikitin, Inna Potekhina, Anastasia Papathanasiou, Dušan Borić, Clive Bonsall, Johannes Krause, Ron Pinhasi, David Reich

**Affiliations:** Department of Genetics, Harvard Medical School, Boston 02115 MA USA; Department of Archaeogenetics, Max Planck Institute for the Science of Human History, 07745 Jena, Germany; Institute for Archaeological Sciences, University of Tuebingen, Germany; Laboratory of Archaeogenetics, Institute of Archaeology, Research Centre for the Humanities, Hungarian Academy of Sciences, H-1097 Budapest, Hungary; Howard Hughes Medical Institute, Harvard Medical School, Boston 02115 MA USA; Earth Institute and School of Archaeology, University College Dublin, Belfield, Dublin 4, Republic of Ireland; Department of Anthropology, University of Vienna, Althanstrasse 14, 1090 Vienna, Austria; CIAS, Department of Life Sciences, University of Coimbra, 3000-456 Coimbra, Portugal; Department of Life Sciences and Biotechnology, University of Ferrara, Via L. Borsari 46. Ferrara 44100 Italy; Australian Centre for Ancient DNA, School of Biological Sciences, The University of Adelaide, SA-5005 Adelaide, Australia; Smurfit Institute of Genetics, Trinity College Dublin, Dublin 2, Ireland; Department of Zoology, University of Cambridge, Downing Street, Cambridge CB2 3EJ, UK; Institute for Anthropological Research, Ljudevita Gaja 32, 10000 Zagreb, Croatia; Broad Institute of Harvard and MIT, Cambridge MA; Department of Anthropology, Emory University, Atlanta, Georgia 30322, USA; Dipartimento di Biologia, Università di Firenze, 50122 Florence, Italy; National Institute of Archaeology and Museum, Bulgarian Academy of Sciences, 2 Saborna Str., BG-1000 Sofia, Bulgaria; Danube Private University, A-3500 Krems, Austria; Department of Biomedical Engineering and Integrative Prehistory and Archaeological Science, CH-4123 Basel-Allschwil, Switzerland; State Office for Heritage Management and Archaeology Saxony-Anhalt and State Museum of Prehistory, D-06114 Halle, Germany; Romanian National History Museum, Bucharest, Romania; Institute of Archaeology, Belgrade, Serbia; Institute of Experimental Morphology, Pathology and Anthropology with Museum, Bulgarian Academy of Sciences, Sofia, Bulgaria; Department of Geosciences, Biogeology, Universität Tübingen, Hölderlinstr. 12, 72074 Tübingen, Germany; Senckenberg Centre for Human Evolution and Palaeoenvironment, University of Tuebingen, 72072 Tuebingen, Germany; Heidelberg Academy of Sciences and Humanities, Research Center ‘‘The Role of Culture in Early Expansions of Humans’’ at the University of Tuebingen, Rümelinstraße 23, 72070 Tuebingen, Germany; ‘Vasile Pârvan’ Institute of Archaeology, Romanian Academy; Human Biology Department, Cardinal Stefan Wyszyński University, Warsaw, Poland; KADUCEJ d.o.o Papandopulova 27, 21000 Split, Croatia; St. Cyril and Methodius University, Veliko Turnovo, Bulgaria; Department of Early Prehistory and Quaternary Ecology, University of Tuebingen, Schloss Hohentübingen, 72070 Tuebingen, Germany; INRAP/UMR 8215 Trajectoires, 21 Alleé de l’Université, 92023 Nanterre, France; Archaeological Museum of Istria, Carrarina 3, 52100 Pula, Croatia; Service Régional de l’Archéologie de Bourgogne-Franche-Comté, 7 rue Charles Nodier, 25043 Besançon Cedex, France; Laboratoire Chronoenvironnement, UMR 6249 du CNRS, UFR des Sciences et Techniques, 16 route de Gray, 25030 Besançon Cedex, France; Regional Museum of History Veliko Tarnovo, Veliko Tarnovo, Bulgaria; Institute for Archaeological Sciences, Paleoanthropology, University of Tuebingen, Rümelinstraße 23, 72070 Tuebingen, Germany; Laboratory for human bio-archaeology, Bulgaria, 1202 Sofia, 42, George Washington str; Regional Museum of History, Vratsa, Bulgaria; DRAC Auvergne – Rhône Alpes, Ministère de la Culture, Le Grenier d’abondance 6, quai Saint Vincent 69283 LYON cedex 01; Eötvös Loránd University, Faculty of Science, Institute of Biology, Department of Biological Anthropology, H-1117 Pázmány Péter sétány 1/c. Budapest, Hungary; Department of Archaeology, Sofia University St. Kliment Ohridski, Bulgaria; Oxford Radiocarbon Accelerator Unit, Research Laboratory for Archaeology and the History of Art, University of Oxford, Dyson Perrins Building, South Parks Road, OX1 3QY Oxford, UK; Regional Museum of History, Haskovo, Bulgaria; Department of Anthropology, University of Wyoming, 1000 E. University Avenue, Laramie, WY 82071, USA; Department of Archaeology, Faculty of Humanities and Social Sciences, University of Zagreb, Ivana Lučića 3, 10000 Zagreb, Croatia; Department of Anthropology and Institutes for Energy and the Environment, Pennsylvania State University, University Park, PA 16802; Department of Bioarchaeology, Institute of Archaeology, National Academy of Sciences of Ukraine; CHU Sainte-Justine Research Center, Pediatric Department, Université de Montréal, Montreal, PQ, Canada, H3T 1C5; National History Museum of Romania, Calea Victoriei, no. 12, 030026, Bucharest, Romania; University of Bucharest, Mihail Kogalniceanu 36-46, 50107, Bucharest, Romania; Institute for Pre- and Protohistoric Archaeology and the Archaeology of the Roman Provinces, Ludwig-Maximilians-University, Schellingstr. 12, 80799 Munich, Germany; Dipartimento SAGAS – Sezione di Archeologia e Antico Oriente, Università degli Studi di Firenze, 50122 Florence, Italy; Museo e Istituto fiorentino di Preistoria, 50122 Florence, Italy; School of History, Classics and Archaeology, University of Edinburgh, Edinburgh EH8 9AG, United Kingdom; Conservation Department in Šibenik, Ministry of Culture of the Republic of Croatia, Jurja Čulinovića 1, 22000 Šibenik, Croatia; Teleorman County Museum, str. 1848, no. 1, 140033 Alexandria, Romania; Peter the Great Museum of Anthropology and Ethnography (Kunstkamera) RAS, 199034 St. Petersburg, Russia; University of Wisconsin, Madison WI, USA; Olga Necrasov Centre for Anthropological Research, Romanian Academy – Iași Branch, Theodor Codrescu St. 2, P.C. 700481, Iași, Romania; Dipartimento di Scienze e tecnologie biologiche, chimiche e farmaceutiche, Lab. of Anthropology, Università degli studi di Palermo, Italy; Anthropological Center, Croatian Academy of Sciences and Arts, 10000 Zagreb, Croatia; Regional Historical Museum Varna, Maria Luiza Blvd. 41, BG-9000 Varna, Bulgaria; National Museum in Belgrade, 1a Republic sq., Belgrade, Serbia; Department of Human Evolution, Max Planck Institute for Evolutionary Anthropology, 04103 Leipzig, Germany; Department of Anthropology, Natural History Museum Vienna, 1010 Vienna, Austria; INRAP/UMR 8215 Trajectoires, 21 Allée de l’Université, 92023 Nanterre, France; CNRS/UMR 7041 ArScAn MAE, 21 Allée de l’Université, 92023 Nanterre, France; Institute of Ethnology and Anthropology, Russian Academy of Sciences, Leninsky Pr., 32a, Moscow, 119991, Russia; Archaeological Museum of Macedonia, Skopje; Regional museum of history, Shumen, Bulgaria; Department of Anthropology, University of Toronto, Toronto, Ontario, M5S 2S2, Canada; Institute of Archaeology & Ethnography, Siberian Branch, Russian Academy of Sciences, Lavrentiev Pr. 17, Novosibirsk 630090, Russia; Satu Mare County Museum Archaeology Department,V. Lucaciu, nr.21, Satu Mare, Romania; Municipal Museum Drniš, Domovinskog rata 54, 22320 Drniš, Croatia; anthropol – Anthropologieservice, Schadenweilerstraße 80, 72379 Hechingen, Germany; Institute for Prehistory, Early History and Medieval Archaeology, University of Tuebingen, Germany; Institute of Latvian History, University of Latvia, Kalpaka Bulvāris 4, Rīga 1050, Latvia; Department of Archaeology, Durham University, UK; School of Environmental Sciences: Geography, University of Hull, Hull HU6 7RX, UK; Department of Biology, Grand Valley State University, Allendale, Michigan, USA; Ephorate of Paleoanthropology and Speleology, Athens, Greece; The Italian Academy for Advanced Studies in America, Columbia University, 1161 Amsterdam Avenue, New York, NY 10027, USA.

## Abstract

Farming was first introduced to southeastern Europe in the mid-7^th^ millennium BCE – brought by migrants from Anatolia who settled in the region before spreading throughout Europe. To clarify the dynamics of the interaction between the first farmers and indigenous hunter-gatherers where they first met, we analyze genome-wide ancient DNA data from 223 individuals who lived in southeastern Europe and surrounding regions between 12,000 and 500 BCE. We document previously uncharacterized genetic structure, showing a West-East cline of ancestry in hunter-gatherers, and show that some Aegean farmers had ancestry from a different lineage than the northwestern Anatolian lineage that formed the overwhelming ancestry of other European farmers. We show that the first farmers of northern and western Europe passed through southeastern Europe with limited admixture with local hunter-gatherers, but that some groups mixed extensively, with relatively sex-balanced admixture compared to the male-biased hunter-gatherer admixture that prevailed later in the North and West. Southeastern Europe continued to be a nexus between East and West after farming arrived, with intermittent genetic contact from the Steppe up to 2,000 years before the migration that replaced much of northern Europe’s population.

## Introduction

The southeastern quadrant of Europe was the beachhead in the spread of agriculture from its source in the Fertile Crescent of southwestern Asia. After the first appearance of agriculture in the mid-7^th^ millennium BCE,^1,2^ farming spread westward via a Mediterranean and northwestward via a Danubian route, and was established in both Iberia and Central Europe by 5600 BCE.^3,4^ Ancient DNA studies have shown that the spread of farming across Europe was accompanied by a massive movement of people^5–8^ closely related to the farmers of northwestern Anatolia^9–11^ but nearly all the ancient DNA from Europe’s first farmers is from central and western Europe, with only three individuals reported from the southeast.^9^ In the millennia following the establishment of agriculture in the Balkan Peninsula, a series of complex societies formed, culminating in sites such as the mid-5^th^ millennium BCE necropolis at Varna, which has some of the earliest evidence of extreme inequality in wealth, with one individual (grave 43) from whom we extracted DNA buried with more gold than is known from any earlier site. By the end of the 6^th^ millennium BCE, agriculture had reached eastern Europe, in the form of the Cucuteni-Trypillian complex in the area of present-day Moldova, Romania and Ukraine, including “mega-sites” that housed hundreds, perhaps thousands, of people.^12^ After around 4000 BCE, these settlements were largely abandoned, and archaeological evidence documents cultural contacts with peoples of the Eurasian steppe.^13^ However, the population movements that accompanied these events have been unknown due to the lack of ancient DNA.

## Results

We generated genome-wide data from 223 ancient humans (214 reported for the first time), from the Balkan Peninsula, the Carpathian Basin, the North Pontic Steppe and neighboring regions, dated to 12,000-500 BCE (Figure 1A, Supplementary Information Table 1, Supplementary Information Note 1). We extracted DNA from skeletal remains in dedicated clean rooms, built DNA libraries and enriched for DNA fragments overlapping 1.24 million single nucleotide polymorphisms (SNPs), then sequenced the product and restricted to libraries with evidence of authentic ancient DNA.^7,10,14^ We filtered out individuals with fewer than 15,000 SNPs covered by at least one sequence, that had unexpected ancestry for their archaeological context and were not directly dated. We report, but do not analyze, nine individuals that were first-degree relatives of others in the dataset, resulting in an analysis dataset of 214 individuals. We analyzed these data together with 274 previously reported 9-11,15-27 ancient individuals,^9–11,15–27^ 799 present-day individuals genotyped on the Illumina “Human Origins” array,^23^ and 300 high coverage genomes from the Simons Genome Diversity Project (SGDP).^28^ We used principal component analysis (PCA; Figure 1B, Extended Data Figure 1), supervised and unsupervised ADMIXTURE (Figure 1D, Extended Data Figure 2),^29^ *D-* statistics, *qpAdm* and *qpGraph,*^30^ along with archaeological and chronological information to cluster the individuals into populations and investigate the relationships among them.

**Figure 1:**
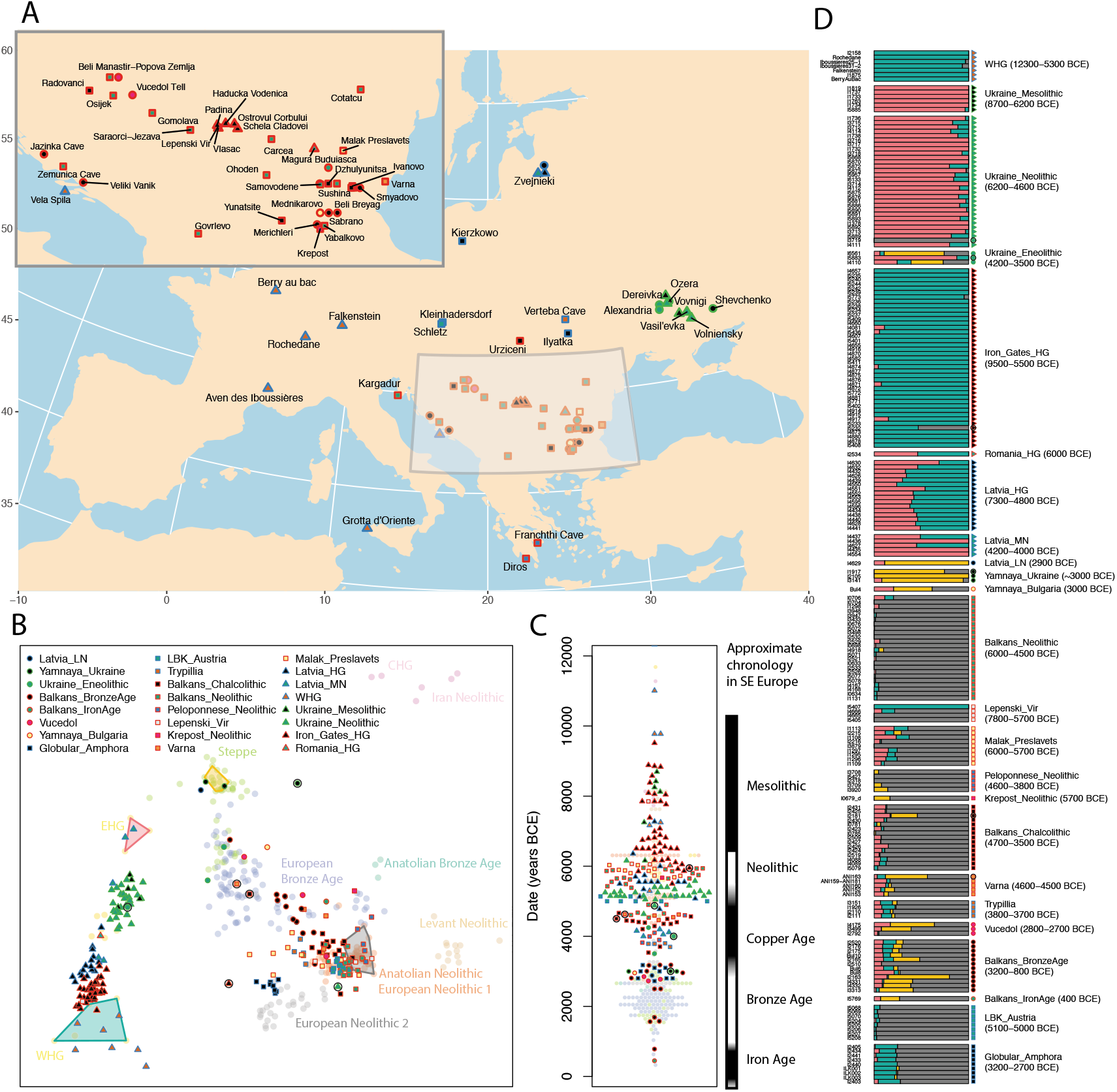
Geographic locations and genetic structure of newly reported individuals. **A**: Location and groupings of newly reported individuals. B: Individuals projected onto axes defined by the principal components of 799 present-day West Eurasians (not shown in this plot for clarity, but shown in Extended Data Figure 1). Projected points include selected published individuals (faded colored circles, labeled) and newly reported individuals (other symbols; outliers shown by additional black circles). Colored polygons indicate the individuals that had cluster memberships fixed at 100% for the supervised admixture analysis in D. C: Estimated age (direct or contextual) for each sample. Approximate chronology used in southeastern Europe shown to the right D: Supervised ADMIXTURE plot, modeling each ancient individual (one per row), as a mixture of populations represented by clusters containing Anatolian Neolithic (grey), Yamnaya from Samara (yellow), EHG (pink) and WHG (green). Dates indicate approximate range of individuals in each population. Map data in A from the *R* package *mapdata*.

We described the individuals in our dataset in terms of their genetic relatedness to a hypothesized set of ancestral populations, which we refer to as their genetic ancestry. It has previously been shown that the great majority of European ancestry derives from three distinct sources.^23^ First, there is “hunter-gatherer-related” ancestry that is more closely related to Mesolithic hunter-gatherers from Europe than to any other population, and that can be further subdivided into “Eastern” (EHG) and “Western” (WHG) hunter-gatherer-related ancestry.^7^ Second, there is “NW Anatolian Neolithic-related” ancestry related to the Neolithic farmers of northwest Anatolia and tightly linked to the appearance of agriculture.^9,10^ The third source, “steppe-related” ancestry, appears in Western Europe during the Late Neolithic to Bronze Age transition and is ultimately derived from a population related to Yamnaya steppe pastoralists.^7,15^ Steppe-related ancestry itself can be modeled as a mixture of EHG-related ancestry, and ancestry related to Upper Palaeolithic hunter-gatherers of the 19,21,22 Caucasus (CHG) and the first farmers of northern Iran.^19,21,22^

### Hunter-Gatherer substructure and transitions

Of the 214 new individuals we report, 114 from Paleolithic, Mesolithic and eastern European Neolithic contexts have almost entirely hunter-gatherer-related ancestry (in eastern Europe, unlike western Europe, “Neolithic” refers to the presence of pottery,^31–33^ not necessarily to farming). These individuals form a cline from WHG to EHG that is correlated with geography (Figure 1B), although it is neither geographically nor temporally uniform (Figure 2, Extended Data Figure 3), and there is also substructure in phenotypically important variants (Supplementary Information Note 2).

**Figure 2:**
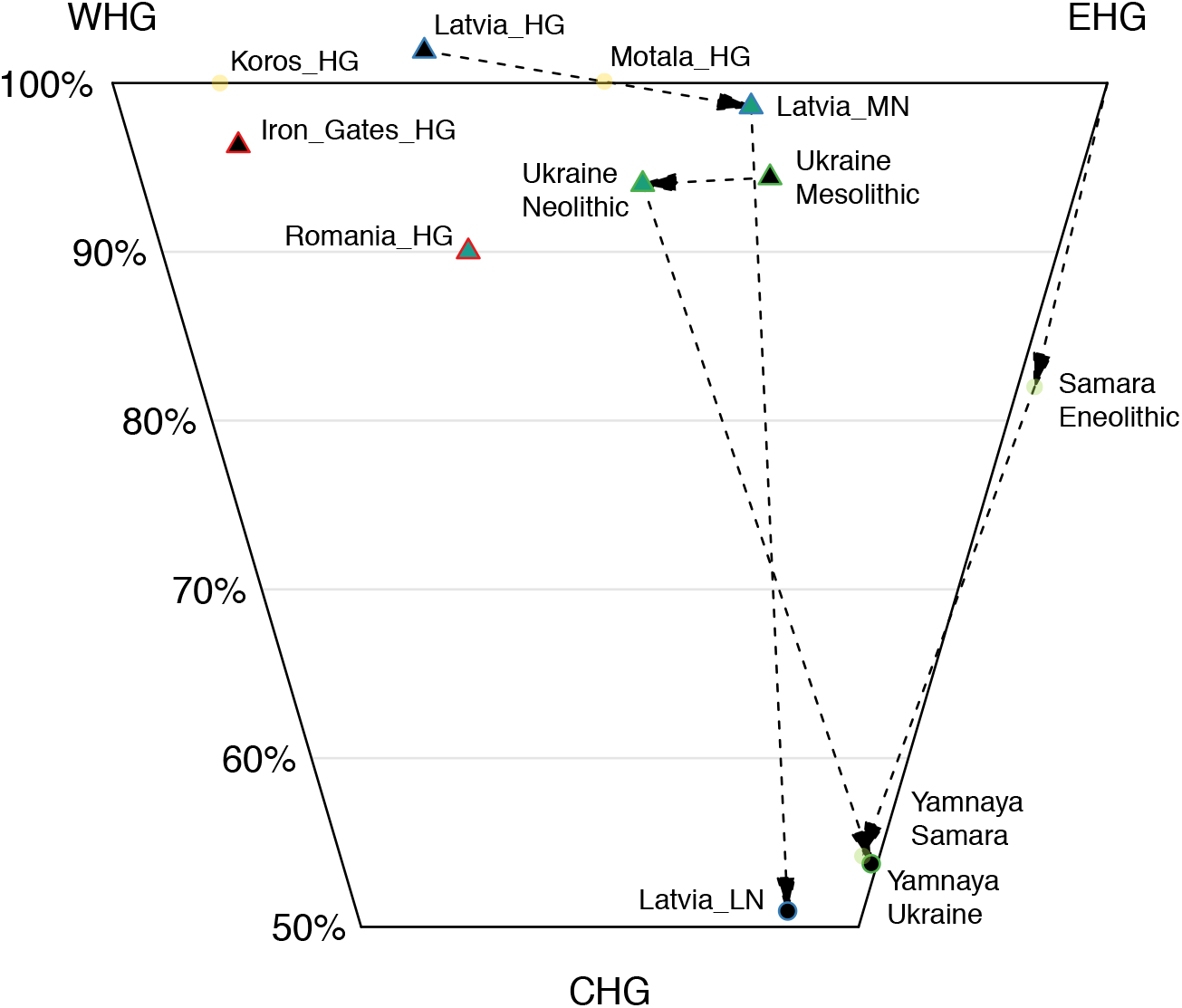
Structure and population change in European populations with hunter-gatherer-related ancestry. This figure shows inferred ancestry proportions for populations modeled as a mixture of WHG, EHG and CHG (Supplementary Table S3.1.3). Dashed lines show populations from the same geographic region. Standard errors range from 1.5-8.3% (Supplementary Table S3.1.3).

From present-day Ukraine, our study reports new genome-wide data from five Mesolithic individuals from ∼9500-6000 BCE, and 31 Neolithic individuals from ∼6000-3500 BCE. On the cline from WHG- to EHG-related ancestry, the Mesolithic individuals fall towards the East, intermediate between EHG and Mesolithic hunter-gatherers from Sweden (Figure 1B).^7^ The Neolithic population has a significant difference in ancestry compared to the Mesolithic (Figures 1B, Figure 2), with a shift towards WHG shown by the statistic D(Mbuti, WHG, Ukraine_Mesolithic, Ukraine_Neolithic); Z=8.9 (Supplementary Information Table 2). Unexpectedly, one Neolithic individual from Dereivka (I3719), which we directly date to 4949-4799 BCE, has entirely NW Anatolian Neolithic-related ancestry.

The pastoralist Bronze Age Yamnaya complex originated on the Eurasian steppe and is a plausible source for the dispersal of steppe-related ancestry into central and western Europe around 2500 BCE.^13^ All previously reported Yamnaya individuals were from Samara^7^ and Kalmykia^15^ in southwest Russia, and had entirely steppe-related ancestry. Here, we report three Yamnaya individuals from further West – from Ukraine and Bulgaria – and show that while they all have high levels of steppe-related ancestry, one from Ozera in Ukraine and one from Bulgaria (I1917 and Bul4, both dated to ∼3000 BCE) have NW Anatolian Neolithic-related admixture, the first evidence of such ancestry in Yamnaya –associated individuals (Figure 1B,D, Supplementary Data Table 2). Two Copper Age individuals (I4110 and I6561, Ukraine_Eneolithic) from Dereivka and Alexandria dated to ∼3600-3400 BCE (and thus preceding the Yamnaya complex) also have mixtures of steppe- and NW Anatolian Neolithic-related ancestry (Figure 1D, Supplementary Data Table 2).

At Zvejnieki in Latvia (17 newly reported individuals, and additional data for 5 first reported in Ref. 34) we observe a transition in hunter-gatherer-related ancestry that is the opposite of that seen in Ukraine. We find (Supplementary Data Table 3) that Mesolithic and Early Neolithic individuals (Latvia_HG) associated with the Kunda and Narva cultures have ancestry intermediate between WHG (∼70%) and EHG (∼30%), consistent with previous reports.^34–36^ We also detect a shift in ancestry between the Early Neolithic and individuals associated with the Middle Neolithic Comb Ware Complex (Latvia_MN), who have more EHG-related ancestry (we estimate 65% EHG, but two of four individuals appear almost 100% EHG in PCA). The most recent individual, associated with the Final Neolithic Corded Ware Complex (I4629, Latvia_LN), attests to another ancestry shift, clustering closely with Yamnaya from Samara,^7^ Kalmykia^15^ and Ukraine (Figure 2).

We report new Upper Palaeolithic and Mesolithic data from southern and western Europe.^17^ Sicilian (I2158) and Croatian (I1875) individuals dating to ∼12,000 and 6100 BCE cluster with previously reported western hunter-gatherers (Figure 1B&D), including individuals from Loschbour^23^ (Luxembourg, 6100 BCE), Bichon^19^ (Switzerland, 11,700 BCE), and Villabruna^17^ (Italy 12,000 BCE). These results demonstrate that WHG populations^23^ were widely distributed from the Atlantic seaboard of Europe in the West, to Sicily in the South, to the Balkan Peninsula in the Southeast, for at least six thousand years.

A particularly important hunter-gatherer population that we report is from the Iron Gates region that straddles the border of present-day Romania and Serbia. This population (Iron_Gates_HG) is represented in our study by 40 individuals from five sites. Modeling Iron Gates hunter-gatherers as a mixture of WHG and EHG (Supplementary Table 3) shows that they are intermediate between WHG (∼85%) and EHG (∼15%). However, this *qpAdm* model does not fit well (p=0.0003, Supplementary table 3) and the Iron Gates hunter-gatherers carry mitochondrial haplogroup K1 (7/40) as well as other subclades of haplogroups U (32/40) and H (1/40). This contrasts with WHG, EHG and Scandinavian hunter-gatherers who almost all carry haplogroups U5 or U2. One interpretation is that the Iron Gates hunter-gatherers have ancestry that is not present in either WHG or EHG. Possible scenarios include genetic contact between the ancestors of the Iron Gates population and Anatolia, or that the Iron Gates population is related to the source population from which the WHG split during a re-expansion into Europe from the Southeast after the Last Glacial Maximum.^17,37^

A notable finding from the Iron Gates concerns the four individuals from the site of Lepenski Vir, two of whom (I4665 & I5405, 6200-5600 BCE), have entirely NW Anatolian Neolithic-related ancestry. Strontium and Nitrogen isotope data^38^ indicate that both these individuals were migrants from outside the Iron Gates, and ate a primarily terrestrial diet (Supplementary Information section 1). A third individual (I4666, 6070 BCE) has a mixture of NW Anatolian Neolithic-related and hunter-gatherer-related ancestry and ate a primarily aquatic diet, while a fourth, probably earlier, individual (I5407) had entirely hunter-gatherer-related ancestry (Figure 1D, Supplementary Information section 1). We also identify one individual from Padina (I5232), dated to 5950 BCE that had a mixture of NW Anatolian Neolithic-related and hunter-gatherer-related ancestry. These results demonstrate that the Iron Gates was a region of interaction between groups distinct in both ancestry and subsistence strategy.

### Population transformations in the first farmers

Neolithic populations from present-day Bulgaria, Croatia, Macedonia, Serbia and Romania cluster closely with the NW Anatolian Neolithic farmers (Figure 1), consistent with archaeological evidence.^39^ Modeling Balkan Neolithic populations as a mixture of NW Anatolian Neolithic and WHG, we estimate that 98% (95% confidence interval [CI]; 97-100%) of their ancestry is NW Anatolian Neolithic-related. A striking exception is evident in 8 out of 9 individuals from Malak Preslavets in present-day Bulgaria.^40^ These individuals lived in the mid-6^th^ millennium BCE and have significantly more hunter-gatherer-related ancestry than other Balkan Neolithic populations (Figure 1B,D, Extended Data Figures 1–3, Supplementary Tables 2-4); a model of 82% (CI: 77-86%) NW Anatolian Neolithic-related, 15% (CI: 12-17%) WHG-related, and 4% (CI: 0-9%) EHG-related ancestry is a fit to the data. This hunter-gatherer-related ancestry with a ∼4:1 WHG:EHG ratio plausibly represents a contribution from local Balkan hunter-gatherers genetically similar to those of the Iron Gates. Late Mesolithic hunter-gatherers in the Balkans were likely concentrated along the coast and major rivers such as the Danube,^41^ which directly connects the Iron Gates with Malak Preslavets. Thus, early farmer groups with the most hunter-gatherer-related ancestry may have been those that lived close to the highest densities of hunter-gatherers.

In the Balkans, Copper Age populations (Balkans_Chalcolithic) harbor significantly more hunter-gatherer-related ancestry than Neolithic populations as shown, for example, by the statistic D(Mbuti, WHG, Balkans_Neolithic, Balkans_Chalcolithic); Z=4.3 ( Supplementary Data Table 2). This is roughly contemporary with the “resurgence” of hunter-gatherer ancestry previously reported in central Europe and Iberia^7,10,42^ and is consistent with changes in funeral rites, specifically the reappearance around 4500 BCE of the Mesolithic tradition of extended supine burial – in contrast to the Early Neolithic tradition of flexed burial.^43^ Four individuals associated with the Copper Age Trypillian population have ∼80% NW Anatolian-related ancestry (Supplementary Table 3), confirming that the ancestry of the first farmers of present-day Ukraine was largely derived from the same source as the farmers of Anatolia and western Europe. Their ∼20% hunter-gatherer ancestry is intermediate between WHG and EHG, consistent with deriving from the Neolithic hunter-gatherers of the region.

We also report the first genetic data associated with the Late Neolithic Globular Amphora Complex. Individuals from two Globular Amphora sites in Poland and Ukraine form a tight cluster, showing high similarity over a large distance (Figure 1B,D). Both Globular Amphora Complex groups of samples had more hunter-gatherer-related ancestry than Middle Neolithic groups from Central Europe^7^ (we estimate 25% [CI: 22-27%] WHG ancestry, similar to Chalcolithic Iberia, Supplementary Data Table 3). In east-central Europe, the Globular Amphora Complex preceded or abutted the Corded Ware Complex that marks the appearance of steppe-related ancestry,^7,15^ while in southeastern Europe, the Globular Amphora Complex bordered populations with steppe-influenced material cultures for hundreds of years^44^ and yet the individuals in our study have no evidence of steppe-related ancestry, providing support for the hypothesis that this material cultural frontier was also a barrier to gene flow.

The movements from the Pontic-Caspian steppe of individuals similar to those associated with the Yamnaya Cultural Complex in the 3^rd^ millennium BCE contributed about 75% of the ancestry of individuals associated with the Corded Ware Complex and about 50% of the ancestry of succeeding material cultures such as the Bell Beaker Complex in central Europe.^7,15^ In two directly dated individuals from southeastern Europe, one (ANI163) from the Varna I cemetery dated to 4711-4550 BCE and one (I2181) from nearby Smyadovo dated to 4550-4450 BCE, we find far earlier evidence of steppe-related ancestry (Figure 1B,D). These findings push back the first evidence of steppe-related ancestry this far West in Europe by almost 2,000 years, but it was sporadic as other Copper Age (∼5000-4000 BCE) individuals from the Balkans have no evidence of it. Bronze Age (∼3400-1100 BCE) individuals do have steppe-related ancestry (we estimate 30%; CI: 26-35%), with the highest proportions in the four latest Balkan Bronze Age individuals in our data (later than ∼1700 BCE) and the least in earlier Bronze Age individuals (3400-2500 BCE; Figure 1D).

### A novel source of ancestry in Neolithic Europe

An important question about the initial spread of farming into Europe is whether the first farmers that brought agriculture to northern Europe and to southern Europe were derived from a single population or instead represent distinct migrations. We confirm that Mediterranean populations, represented in our study by individuals associated with the Epicardial Early Neolithic from Iberia^7^, are closely related to Danubian populations represented by the *Linearbandkeramik* (LBK) from central Europe^7,45^ and that both are closely related to the Balkan Neolithic population. These three populations form a clade with the NW Anatolian Neolithic individuals as an outgroup, consistent with a single migration into the Balkan peninsula, which then split into two (Supplementary Information Note 3).

In contrast, five southern Greek Neolithic individuals (Peloponnese_Neolithic) – three (plus one previously published^26^) from Diros Cave and one from Franchthi Cave – are not consistent with descending from the same source population as other European farmers. *D*-statistics (Supplementary Information Table 2) show that in fact, these “Peloponnese Neolithic” individuals dated to ∼4000 BCE are shifted away from WHG and towards CHG, relative to Anatolian and Balkan Neolithic individuals. We see the same pattern in a single Neolithic individual from Krepost in present-day Bulgaria (I0679_d, 5718-5626 BCE). An even more dramatic shift towards CHG has been observed in individuals associated with the Bronze Age Minoan and Mycenaean cultures,^26^ and thus there was gene flow into the region from populations with CHG-rich ancestry throughout the Neolithic, Chalcolithic and Bronze Age. Possible sources are related to the Neolithic population from the central Anatolian site of Tepecik Ciftlik,^21^ or the Aegean site of Kumtepe,^11^ who are also shifted towards CHG relative to NW Anatolian Neolithic samples, as are later Copper and Bronze Age Anatolians.^10,26^

### Sex-biased admixture between hunter-gatherers and farmers

We provide the first evidence for sex-biased admixture between hunter-gatherers and farmers in Europe, showing that the Middle Neolithic “resurgence” of hunter-gatherer-related ancestry^7,42^ in central Europe and Iberia was driven more by males than by females (Figure 3B&C, Supplementary Data Table 5, Extended Data Figure 4). To document this we used *qpAdm* to compute ancestry proportions on the autosomes and the X chromosome; since males always inherit their X chromosome from their mothers, differences imply sex-biased mixture. In the Balkan Neolithic there is no evidence of sex bias (Z=0.27 where a positive Z-score implies male hunter-gatherer bias), nor in the LBK and Iberian_Early Neolithic (Z=-0.22 and 0.74). In the Copper Age there is clear bias: weak in the Balkans (Z=1.66), but stronger in Iberia (Z=3.08) and Central Europe (Z=2.74). Consistent with this, hunter-gatherer mitochondrial haplogroups (haplogroup U)^46^ are rare and within the intervals of genome-wide ancestry proportions, but hunter-gatherer-associated Y chromosomes (haplogroups I, R1 and C1)^17^ are more common: 7/9 in the Iberian Neolithic/Copper Age and 9/10 in Middle-Late Neolithic Central Europe (Central_MN and Globular_Amphora) (Figure 3C).

**Figure 3:**
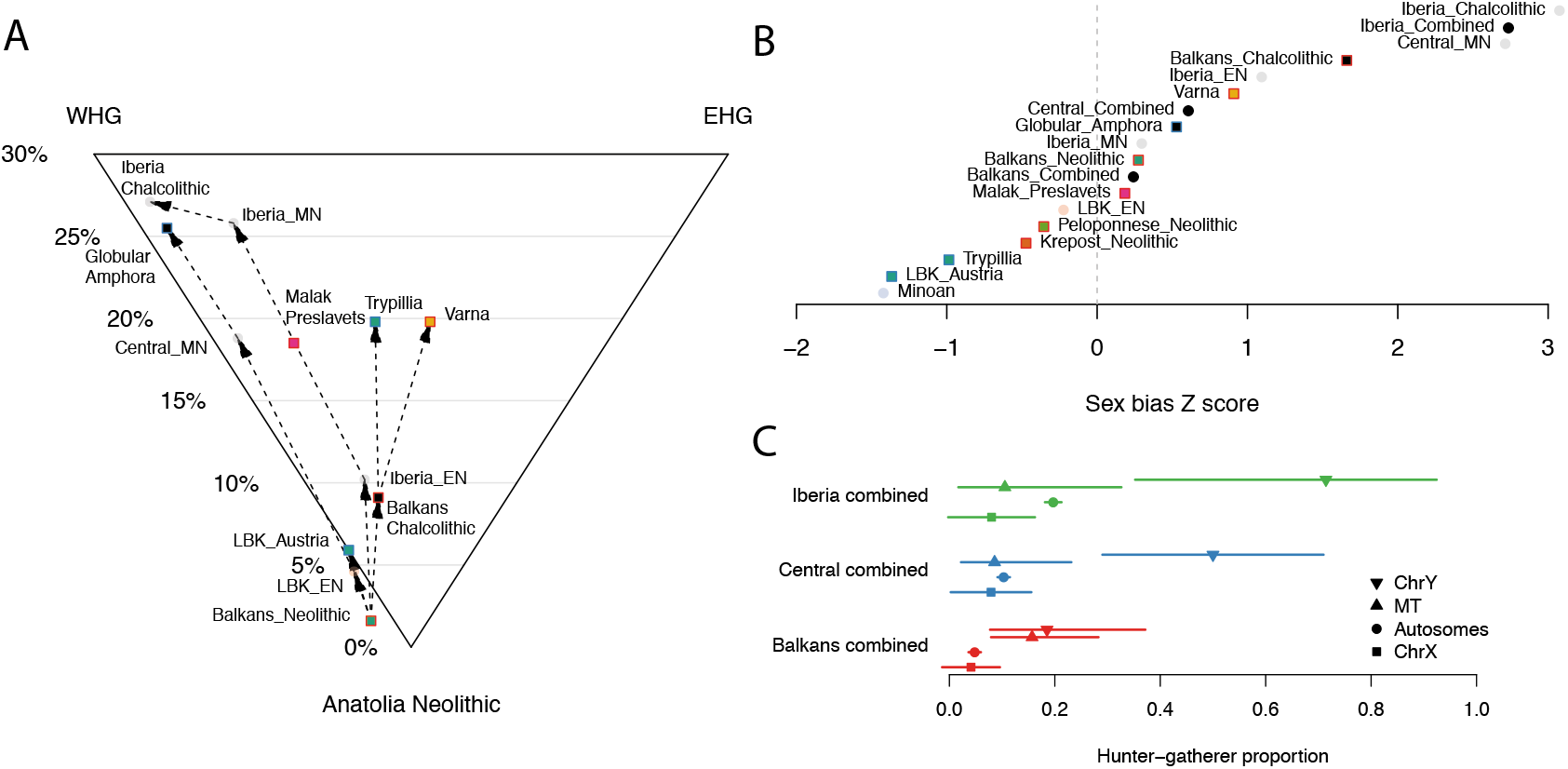
Structure and population change in European populations with NW Anatolian Neolithic-related ancestry. A: each population is modeled as a mixture of NW Anatolia Neolithic, WHG, and EHG. Dashed lines show temporal relationships between populations from the same geographic areas with similar ancestries. Standard errors range from 0.7-6.0% (Supplementary Table S3.2.2). B: Z-scores for the difference in hunter-gatherer-related ancestry on the autosomes compared to the X chromosome when populations are modeled as a mixture of NW Anatolia Neolithic and WHG. A positive score indicates that there is more hunter-gatherer-related ancestry on the autosomes and therefore the hunter-gatherer-related ancestry is male-biased. C: Hunter-gatherer-related ancestry proportions on the autosomes, X chromosome, mitochondrial DNA (i.e. mt haplogroup U), and the Y chromosome (i.e. Y chromosome haplogroups I2, R1 and C2). Bars show approximate 95% confidence intervals. “Combined” populations merge all individuals from different times from a geographic area.

### No evidence that steppe-related ancestry moved through southeast Europe into Anatolia

One version of the Steppe Hypothesis of Indo-European language origins suggests that Proto-Indo-European languages developed north of the Black and Caspian seas, and that the earliest known diverging branch – Anatolian – was spread into Asia Minor by movements of steppe peoples through the Balkan peninsula during the Copper Age around 4000 BCE.^47^ If this were correct, then one way to detect evidence of it would be the appearance of large amounts of steppe-related ancestry first in the Balkan Peninsula, and then in Anatolia. However, our data show no evidence for this scenario. While we find sporadic examples of steppe-related ancestry in Balkan Copper and Bronze Age individuals, this ancestry is rare until the late Bronze Age. Moreover, while Bronze Age Anatolian individuals have CHG-related ancestry,^26^ they have neither the EHG-related ancestry characteristic of all steppe populations sampled to date,^19^ nor the WHG-related ancestry that is ubiquitous in Neolithic southeastern Europe (Extended Data Figure 2, Supplementary Data Table 2). An alternative hypothesis is that the ultimate homeland of Proto-Indo-European languages was in the Caucasus or in Iran. In this scenario, westward movement contributed to the dispersal of Anatolian languages, and northward movement and mixture with EHG was responsible for the formation of a “Late Proto-Indo European”-speaking population associated with the Yamnaya Complex.^13^ While this scenario gains plausibility from our results, it remains possible that Indo-European languages were spread through southeastern Europe into Anatolia without large-scale population movement or admixture.

## Discussion

Our study shows that southeastern Europe consistently served as a genetic contact zone. Before the arrival of farming, the region saw interaction between diverged groups of hunter-gatherers, and this interaction continued after farming arrived. While this study has clarified the genomic history of southeastern Europe from the Mesolithic to the Bronze Age, the processes that connected these populations to the ones living today remain largely unknown. An important direction for future research will be to sample populations from the Bronze Age, Iron Age, Roman, and Medieval periods and to compare them to present-day populations to understand how these transitions occurred.

## Methods

### Ancient DNA Analysis

We extracted DNA and prepared next-generation sequencing libraries in four different dedicated ancient DNA laboratories (Adelaide, Boston, Budapest, and Tuebingen). We also prepared samples for extraction in a fifth laboratory (Dublin), from whence it was sent to Boston for DNA extraction and library preparation (Supplementary Table 1).

Two samples were processed at the Australian Centre for Ancient DNA, Adelaide, Australia, according to previously published methods^7^ and sent to Boston for subsequent screening, 1240k capture and sequencing.

Seven samples were processed^27^ at the Institute of Archaeology RCH HAS, Budapest, Hungary, and amplified libraries were sent to Boston for screening, 1240k capture and sequencing.

Seventeen samples were processed at the Institute for Archaeological Sciences of the University of Tuebingen and at the Max Planck Institute for the Science of Human History in Jena, Germany. Extraction^48^ and library preparation^49,50^ followed established protocols. We performed in-solution capture as described below (“1240k capture”) and sequenced on an Illumina HiSeq 4000 or NextSeq 500 for 76bp using either single- or paired-end sequencing.

The remaining 197 samples were processed at Harvard Medical School, Boston, USA. From about 75mg of sample powder from each sample (extracted in Boston or University College Dublin, Dublin, Ireland), we extracted DNA following established methods^48^ replacing the column assembly with the column extenders from a Roche kit.^51^ We prepared double barcoded libraries with truncated adapters from between one ninth and one third of the DNA extract. Most libraries included in the nuclear genome analysis (90%) were subjected to partial (“half”) Uracil-DNA-glycosylase (UDG) treatment before blunt end repair. This treatment reduces by an order of magnitude the characteristic cytosine-to-thymine errors of ancient DNA data^52^, but works inefficiently at the 5’ ends,^50^ thereby leaving a signal of characteristic damage at the terminal ends of ancient sequences. Some libraries were not UDG-treated (“minus”). For some samples we increased coverage by preparing additional libraries from the existing DNA extract using the partial UDG library preparation, but replacing the MinElute column cleanups in between enzymatic reactions with magnetic bead cleanups, and the final PCR cleanup with SPRI bead cleanup.^53,54^

We screened all libraries from Adelaide, Boston and Budapest by enriching for the mitochondrial genome plus about 3,000 (50 in an earlier, unpublished, version) nuclear SNPs using a bead-capture^55^ but with the probes replaced by amplified oligonucleotides synthesized by CustomArray Inc. After the capture, we completed the adapter sites using PCR, attaching dual index combinations^56^ to each enriched library. We sequenced the products of between 100 and 200 libraries together with the non-enriched libraries (shotgun) on an Illumina NextSeq500 using v2 150 cycle kits for 2x76 cycles and 2x7 cycles.

In Boston, we performed two rounds of in-solution enrichment (“1240k capture”) for a targeted set of 1,237,207 SNPs using previously reported protocols.^7,14,23^ For a total of 34 individuals, we increased coverage by building one to eight additional libraries for the same sample. When we built multiple libraries from the same extract, we often pooled them in equimolar ratios before the capture. We performed all sequencing on an Illumina NextSeq500 using v2 150 cycle kits for 2x76 cycles and 2x7 cycles. We attempted to sequence each enriched library up to the point where we estimated that it was economically inefficient to sequence further. Specifically, we iteratively sequenced more and more from each individual and only stopped when we estimated that the expected increase in the number of targeted SNPs hit at least once would be less than about one for every 100 new read pairs generated. After sequencing, we trimmed two bases from the end of each read and aligned to the human genome (b37/hg19) using *bwa*.^57^ We then removed individuals with evidence of contamination based on mitochondrial DNA polymorphism^58^ or difference in PCA space between damaged and undamaged reads^59^, a high rate of heterozygosity on chromosome X despite being male^59,60^, or an atypical ratio of X-to-Y sequences. We also removed individuals that had low coverage (fewer than 15,000 SNPs hit on the autosomes). We report, but do not analyze, data from nine individuals that were first-degree relatives of others in the dataset (determined by comparing rates of allele sharing between pairs of individuals).

After removing a small number of sites that failed to capture, we were left with a total of 1,233,013 sites of which 32,670 were on chromosome X and 49,704 were on chromosome Y, with a median coverage at targeted SNPs on the 214 newly reported individuals of 0.90 (range 0.007-9.2; Supplementary Table 1). We generated “pseudo-haploid” calls by selecting a single read randomly for each individual at each SNP. Thus, there is only a single allele from each individual at each site, but adjacent alleles might come from either of the two haplotypes of the individual. We merged the newly reported data with previously reported 9-11,15-27 data from 274 other ancient individuals^9–11,15–27^, making pseudo-haploid calls in the same way at the 1240k sites for individuals that were shotgun sequenced rather than captured. Using the captured mitochondrial sequence from the screening process, we called mitochondrial haplotypes. Using the captured SNPs on the Y chromosome, we called Y chromosome haplogroups for males by restricting to sequences with mapping quality ≥30 and bases with base quality ≥30. We determined the most derived mutation for each individual, using the nomenclature of the International Society of Genetic Genealogy (http://www.isogg.org) version 11.110 (21 April 2016).

### Population genetic analysis

To analyze these ancient individuals in the context of present day genetic diversity, we merged them with the following two datasets:

1. 300 high coverage genomes from a diverse worldwide set of 142 populations sequenced as part of the Simons Genome Diversity Project^28^ (SGDP merge).
2. 799 West Eurasian individuals genotyped on the Human Origins array^23^, with 597,573 sites in the merged dataset (HO merge).

We computed principal components of the present-day individuals in the HO merge and projected the ancient individuals onto the first two components using the “*lsqproject: YES*” option in *smartpca (v15100)*^61^ (https://www.hsph.harvard.edu/alkes-price/software/).

We ran *ADMIXTURE (v1.3.0)* in both supervised and unsupervised mode. In supervised mode we used only the ancient individuals, on the full set of SNPs, and the following population labels fixed:

- *Anatolia_Neolithic*
- *WHG*
- *EHG*
- *Yamnaya*

For unsupervised mode we used the HO merge, including 799 present-day individuals. We flagged individuals that were genetic outliers based on PCA and ADMIXTURE, relative to other individuals from the same time period and archaeological culture.

We computed *D-*statistics using *qpDstat (v710)*. *D*-statistics of the form D(A,B,X,Y) test the null hypothesis of the unrooted tree topology ((A,B),(X,Y)). A positive value indicates that either A and X, or B and Y, share more drift than expected under the null hypothesis. We quote *D-*statistics as the *Z*-score computed using default block jackknife parameters.

We fitted admixture proportions with *qpAdm (v610)* using the SGDP merge. Given a set of outgroup (“right”) populations, *qpAdm* models one of a set of source (“left”) populations (the “test” population) as a mixture of the other sources by fitting admixture proportions to match the observed matrix of *f4-*statistics as closely as possible. We report a p-value for the null hypothesis that the test population does not have ancestry from another source that is differentially related to the right populations. We computed standard errors for the mixture proportions using a block jackknife. Importantly, *qpAdm* does not require that the source populations are actually the admixing populations, only that they are a clade with the correct admixing populations, relative to the other sources. Infeasible coefficient estimates (i.e. outside [0,1]) are usually a sign of poor model fit, but in the case where the source with a negative coefficient is itself admixed, could be interpreted as implying that the true source is a population with different admixture proportions. We used the following set of seven populations as outgroups or “right populations”:

- *Mbuti.DG*
- *Ust_Ishim_HG_published.DG*
- *Mota.SG*
- *MA1_HG.SG*
- *Villabruna*
- *Papuan.DG*
- *Onge.DG*
- *Han.DG*

For some analyses where we required extra resolution (Extended Data Table 4) we used an extended set of 14 right (outgroup) populations, including additional Upper Paleolithic European individuals^17^:

- *ElMiron*
- *Mota.SG*
- *Mbuti.DG*
- *Ust_Ishim_HG_published.DG*
- *MA1_HG.SG*
- *AfontovaGora3*
- *GoyetQ116-1_published*
- *Villabruna*
- *Kostenki14*
- *Vestonice16*
- *Karitiana.DG*
- *Papuan.DG*
- *Onge.DG*
- *Han.DG*

We also fitted admixture graphs with *qpGraph (v6021)*^30^ (https://github.com/DReichLab/AdmixTools, Supplementary Information, section 3). Like *qpAdm*, *qpGraph* also tries to match a matrix of *f-*statistics, but rather than fitting one population as a mixture of other, specified, populations, it fits the relationship between all tested populations simultaneously, potentially incorporating multiple admixture events. However, *qpGraph* requires the graph relating populations to be specified in advance. We tested goodness-of-fit by computing the expected *D-*statistics under the fitted model, finding the largest *D*-statistic outlier between the fitted and observed model, and computing a *Z-*score using a block jackknife.

For 116 individuals with hunter-gatherer-related ancestry we estimated an effective migration surface using the software *EEMS* (https://github.com/dipetkov/eems)^62^. We computed pairwise differences between individuals using the *bed2diffs2* program provided with *EEMS*. We set the number of demes to 400 and defined the outer boundary of the region by the polygon (in latitude-longitude co-ordinates) [(66,60), (60,10), (45,-15), (35,-10), (35,60)]. We ran the MCMC ten times with different random seeds, each time with one million burn-in and four million regular iterations, thinned to one in ten thousand.

To analyze potential sex bias in admixture, we used *qpAdm* to estimate admixture proportions on the autosomes (default option) and on the X chromosome (option “*chrom: 23*”). We computed Z-scores for the difference between the autosomes and the X chromosome as = 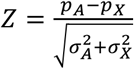 where *p*_*A*_ and *p*_*X*_ are the hunter-gatherer admixture proportions on the autosomes and the X chromosome, and *σ_A_* and *σ_X_* are the corresponding jackknife standard deviations. Thus, a positive Z-score means that there is more hunter-gatherer admixture on the autosomes than on the X chromosome, indicating that the hunter-gatherer admixture was male-biased. Because X chromosome standard errors are high and *qpAdm* results can be sensitive to which population is first in the list of outgroup populations, we checked that the patterns we observe were robust to cyclic permutation of the outgroups. To compare frequencies of hunter-gatherer uniparental markers, we counted the individuals with mitochondrial haplogroup U and Y chromosome haplogroups C2, I2 and R1, which are all common in Mesolithic hunter-gatherers but rare or absent in Anatolian Neolithic individuals. The Iron Gates hunter-gatherers also carry H and K1 mitochondrial haplogroups so the proportion of haplogroup U represents the minimum maternal hunter-gatherer contribution. We computed binomial confidence intervals for the proportion of haplogroups associated with each ancestry type using the Agresti-Coull method^63,64^ implemented in the *binom* package in *R*.

Given autosomal and X chromosome admixture proportions, we estimated the proportion of male and female hunter-gatherer ancestors by assuming a single-pulse model of admixture. If the proportions of male and female ancestors that are hunter-gatherer-related are given by *m* and *f*, respectively, then the proportions of hunter-gatherer-related ancestry on the autosomes and the X chromosome are given by 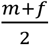 and 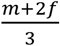. We approximated the sampling error in the observed admixture proportions by the estimated jackknife error and computed the likelihood surface for *(m,f)* over a grid ranging from (0,0) to (1,1).

### Direct AMS ^14^C Bone Dates

We report 113 new direct AMS ^14^C bone dates for 112 individuals from multiple AMS radiocarbon laboratories. In general, bone samples were manually cleaned and demineralized in weak HCl and, in most cases (PSU, UCIAMS, OxA), soaked in an alkali bath (NaOH) at room temperature to remove contaminating soil humates. Samples were then rinsed to neutrality in Nanopure H2O and gelatinized in HCL.^65^ The resulting gelatin was lyophilized and weighed to determine percent yield as a measure of collagen preservation (% crude gelatin yield). Collagen was then directly AMS ^14^C dated (Beta, AA) or further purified using ultrafiltration (PSU, UCIAMS, OxA, Poz, MAMS).^66^ It is standard in some laboratories (PSU/UCIAMS, OxA) to use stable carbon and nitrogen isotopes as an additional quality control measure. For these samples, the %C, %N and C:N ratios were evaluated before AMS ^14^C dating.^67^ C:N ratios for well-preserved samples fall between 2.9 and 3.6, indicating good collagen preservation.^68^ For 94 new samples, we also report δ^13^C and δ^15^N values (Supplementary Table 6).

All ^14^C ages were δ^13^C-corrected for mass dependent fractionation with measured ^13^C/^12^C values^69^ and calibrated with OxCal version 4.2.3^70^ using the IntCal13 northern hemisphere calibration curve.^70^ For hunter-gatherers from the Iron Gates, the direct ^14^C dates tend to be overestimates because of the freshwater reservoir effect (FRE), which arises because of a diet including fish that consumed ancient carbon, and for these individuals we performed a correction (Supplementary Information Note 1),^71^ assuming that 100% FRE = 545±70 yr, and δ^15^N values of 8.3% and 17.0% for 100% terrestrial and aquatic diets, respectively.

## Acknowledgments

We thank David Anthony, Iosif Lazaridis, and Mark Lipson for comments on the manuscript, Bastien Llamas and Alan Cooper for contributions to laboratory work, Richard Evershed for contributing ^14^C dates and Friederike Novotny for assistance with samples. Support for this project was provided by the Human Frontier Science Program fellowship LT001095/2014-L to I.M.; by DFG grant AL 287 / 14-1 to K.W.A.; by Irish Research Council grant GOIPG/2013/36 to D.F.; by the NSF Archaeometry program BCS-1460369 to DJK (for AMS ^14^C work at Penn State); by MEN-UEFISCDI grant, Partnerships in Priority Areas Program – PN II (PN-II-PT-PCCA-2013-4-2302) to C.L.; by Croatian Science Foundation grant IP-2016-06-1450 to M.N.; by European Research Council grant ERC StG 283503 and Deutsche Forschungsgemeinschaft DFG FOR2237 to K.H.; by ERC starting grant ADNABIOARC (263441) to R.P.; and by US National Science Foundation HOMINID grant BCS-1032255, US National Institutes of Health grant GM100233, and the Howard Hughes Medical Institute to D.R.

## Author Contributions

SAR, AS-N, SVai, SA, KWA, RA, DA, AA, NA, KB, MBG, HB, MB, ABo, YB, ABu, JB, SC, NC, RC, MC, CC, DD, NE, MFr, BGal, GG, BGe, THa, VH, KH, THi, SI, IJ, IKa, DKa, AK, DLa, MLa, CL, MLe, KL, DLV, DLo, IL, MMa, FM, KM, HM, MMe, PM, VM, VP, TDP, ASi, LS, MŠ, VS, PS, ASt, TS, MT-N, CT, IV, FVa, SVas, FVe, SV, EV, BV, CV, JZ, SZ, PWS, GC, RK, DC, GZ, BGay, MLi, AGN, IP, AP, DB, CB, JK, RP & DR assembled and interpreted archaeological material. CP, AS-N, NR, NB, FC, OC, DF, MFe, BGam, GGF, WH, EH, EJ, DKe, BK-K, IKu, MMi, AM, KN, MN, JO, SP, KSi, KSt & SVai performed laboratory work. IM, CP, AS-N, SM, IO, NP & DR analyzed data. DJK, ST, DB, CB interpreted ^14^C dates. JK, RP & DR supervised analysis or laboratory work. IM & DR wrote the paper, with input from all co-authors.

## Extended Data Figures

**Extended Data Figure 1:**
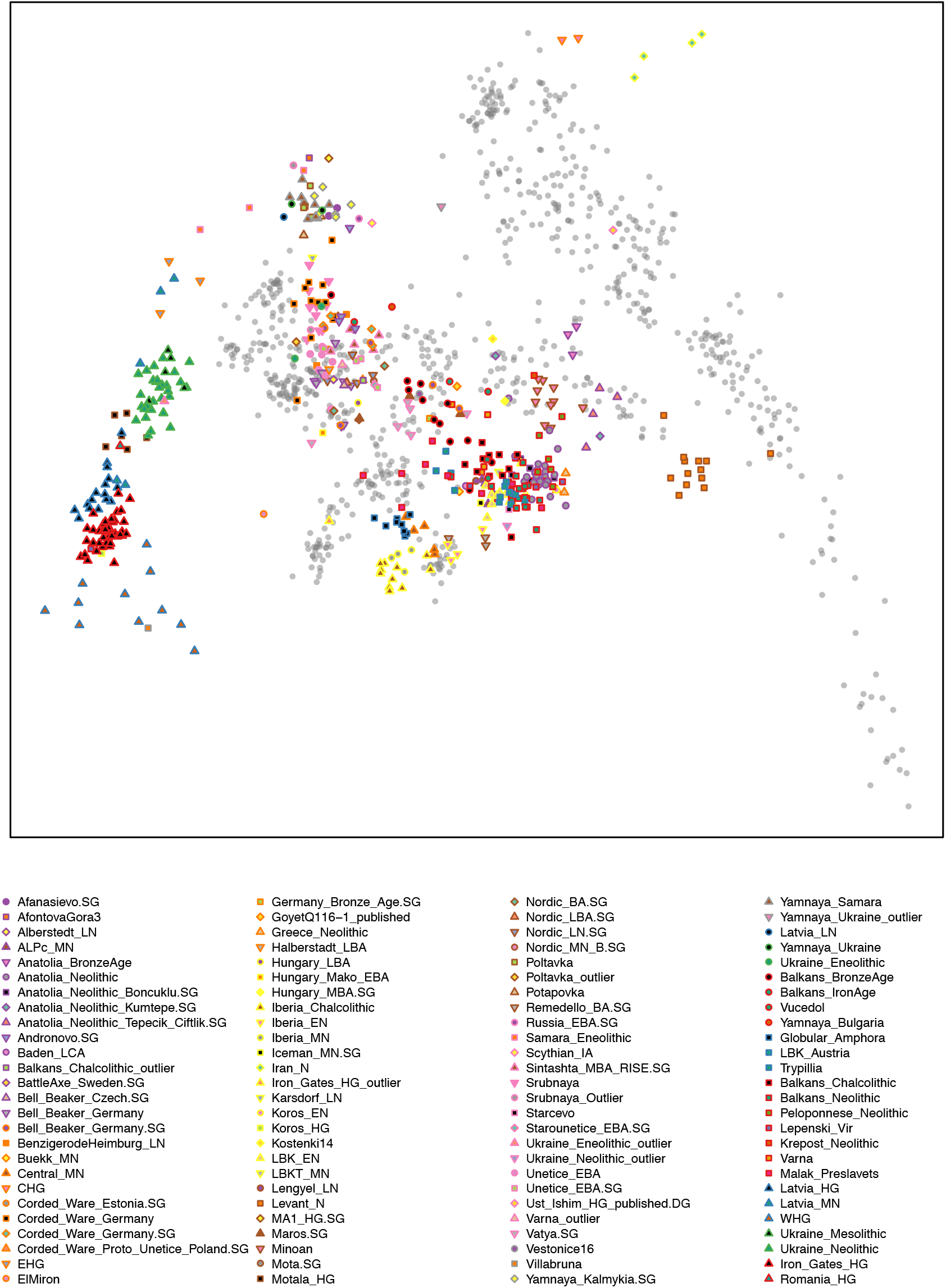
PCA of all ancient individuals, projected onto principal components defined by 799 present-day West Eurasian individuals. (This differs from Figure 1B in that the plot is not cropped and the present-day individuals are shown.)

**Extended Data Figure 2:**
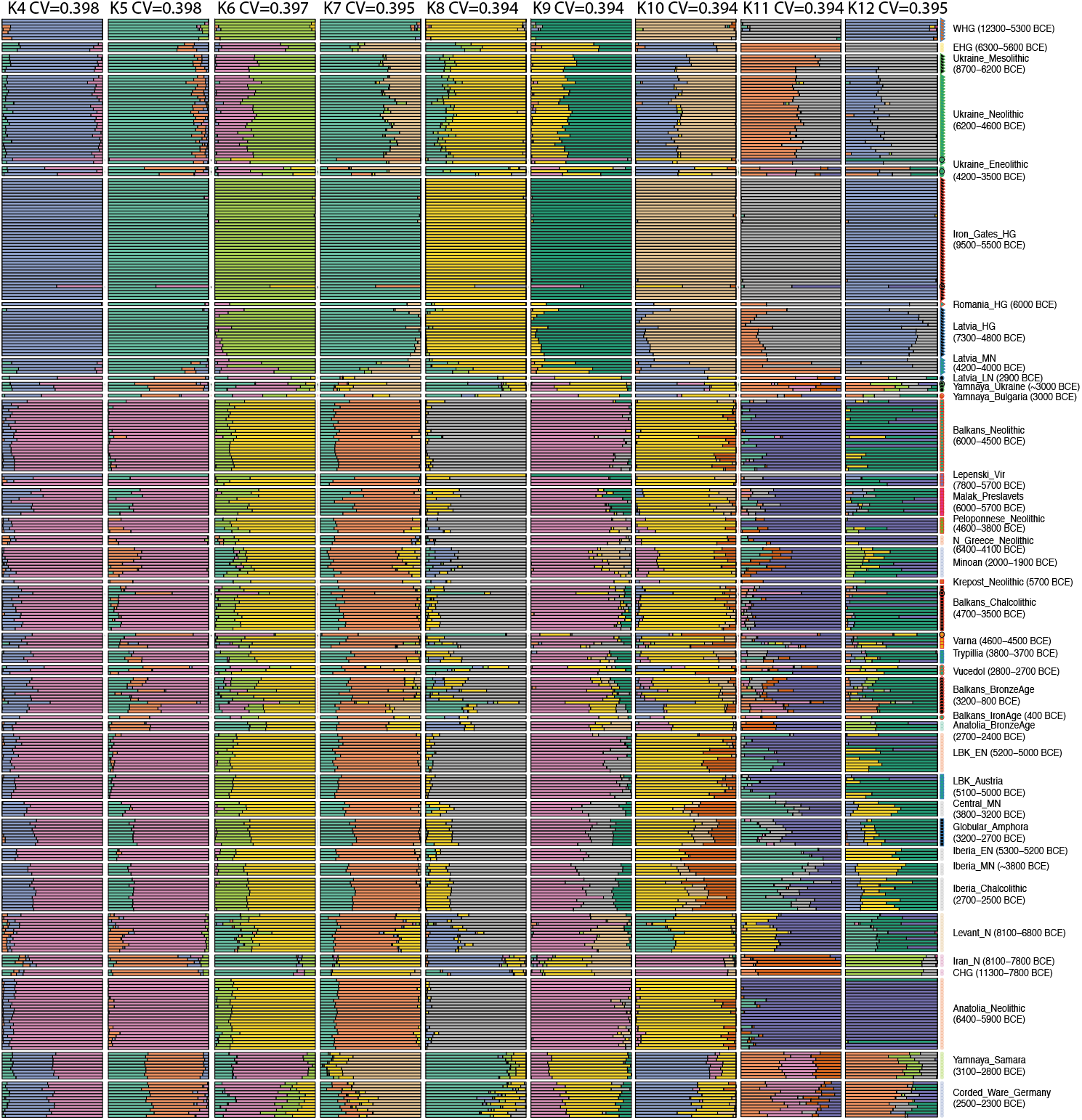
Unsupervised ADMIXTURE plot from k=4 to 12, on a dataset consisting of 1099 present-day individuals and 476 ancient individuals. We show newly reported ancient individuals and some previously published individuals for comparison.

**Extended Data Figure 3:**
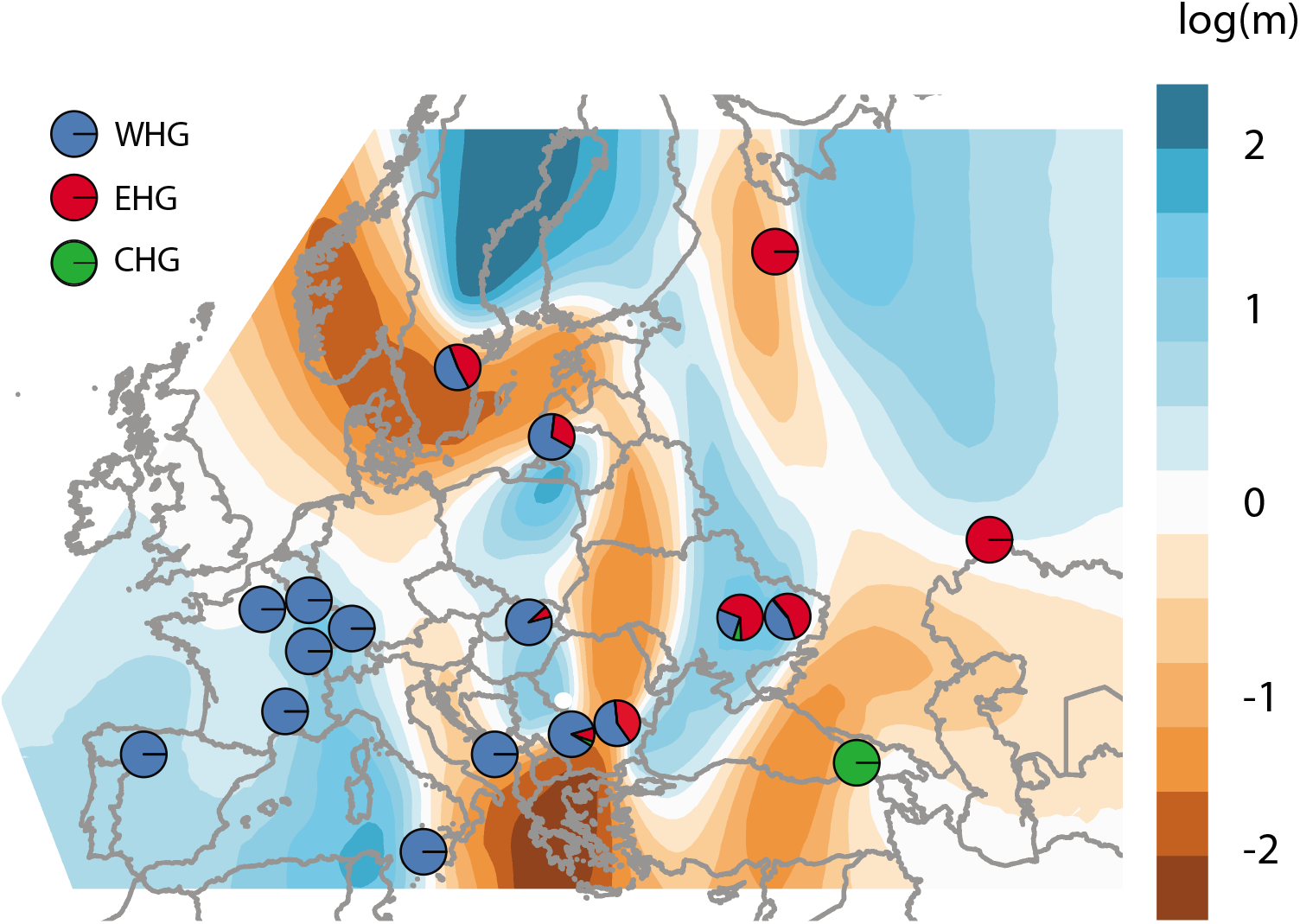
Spatial structure in hunter-gatherers. Estimated effective migration surface (EEMS).^62^ This fits a model of genetic relatedness where individuals move (in a random direction) from generation to generation on an underlying grid so that genetic relatedness is determined by distance. The migration parameter *m* defines the local rate of migration, varies on the grid and is inferred. This plot shows *log10* m, scaled relative to the average migration rate (which is arbitrary). Thus *log10(m)=2*, for example, implies that the rate of migration at this point on the grid is 100 times higher than average. To restrict as much as possible to hunter-gatherer structure, the migration surface is inferred using data from 116 individuals from populations that date earlier than ∼5000 BCE and have no NW Anatolian-related ancestry. Though the migration surface is sensitive to sampling, and fine-scale features may not be interpretable, the migration “barrier” (region of low migration) running north-south and separating populations with primarily WHG from primarily EHG ancestry seems to be robust, and consistent with inferred admixture proportions. This analysis suggests that Mesolithic hunter-gatherer population structure was clustered and not smoothly clinal, in the sense that genetic differentiation did not vary consistently with distance. Superimposed on this background, pies show the WHG, EHG and CHG ancestry proportions inferred for populations used to construct the migration surface (another way of visualizing the data in show in Figure 2, Supplementary Table 3.1.3 – we use two population models if they fit with p>0.1, and three population models otherwise). Pies with only a single color have been fixed to be the source populations.

**Extended Data Figure 4:**
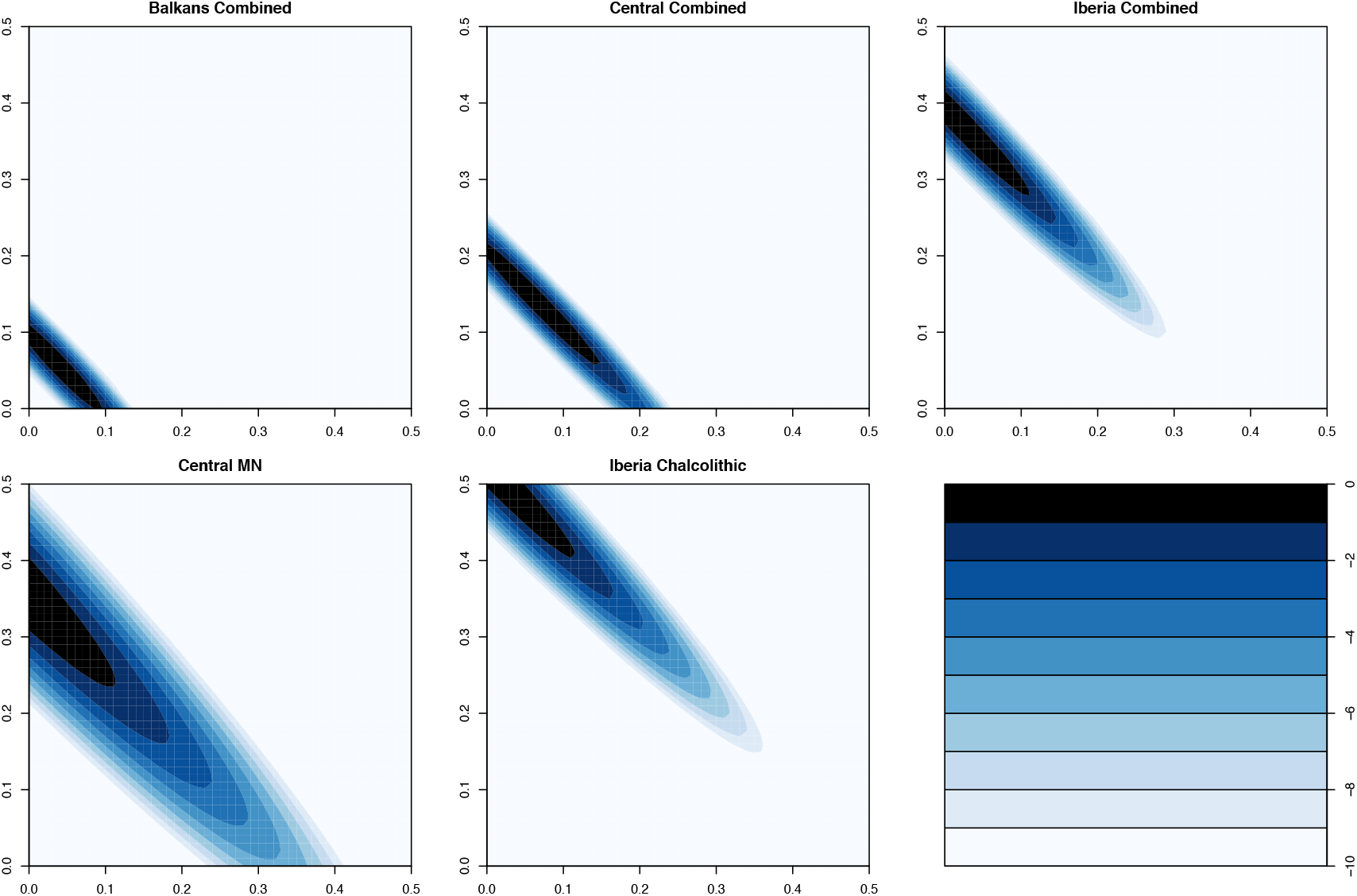
log-likelihood surfaces for the proportion of female (x-axis) and male (y-axis) ancestors that are hunter-gatherer-related for the combined populations analyzed in Figure 3C, and the two populations with the strongest evidence for sex-bias. Log-likelihood scale ranges from 0 to -10, where 0 is the feasible point with the highest likelihood.

## Supplementary Tables

**Supplementary Table 1**: Details of ancient individuals analyzed in this study.

**Supplementary Table 2**: Key *D-*statistics to support statements about population history.

**Supplementary Table 3**: *qpAdm* models with 7-population outgroup set.

**Supplementary Table 4**: *qpAdm* models with extended 14-population outgroup set.

**Supplementary Table 5**: *qpAdm* models for Neolithic populations for chromosome X.

**Supplementary Table 6**: Additional ^14^C dating information.

